# pyBedGraph: a Python package for fast operations on 1-dimensional genomic signal tracks

**DOI:** 10.1101/709683

**Authors:** Henry B. Zhang, Minji Kim, Jeffrey H. Chuang, Yijun Ruan

## Abstract

**Motivation:** Modern genomic research relies heavily on next-generation sequencing experiments such as ChIP-seq and ChIA-PET that generate coverage files for transcription factor binding, as well as DHS and ATAC-seq that yield coverage files for chromatin accessibility. Such files are in a bedGraph text format or a bigWig binary format. Obtaining summary statistics in a given region is a fundamental task in analyzing protein binding intensity or chromatin accessibility. However, the existing Python package for operating on coverage files is not optimized for speed.

**Results:** We developed pyBedGraph, a Python package to quickly obtain summary statistics for a given interval in a bedGraph file. When tested on 8 ChIP-seq and ATAC-seq datasets, pyBedGraph is on average 245 times faster than the existing program. Notably, pyBedGraph can look up the exact mean signal of 1 million regions in ~0.26 second on a conventional laptop. An approximate mean for 10,000 regions can be computed in ~0.0012 second with an error rate of less than 5 percent.

**Availability:** pyBedGraph is publicly available at https://github.com/TheJacksonLaboratory/pyBedGraph under the MIT license.

## 1 Introduction

The advancement of next-generation sequencing technologies allowed researchers to measure various biological signals in the genome. For example, one can probe gene expression (RNA-seq) (Mortazavi et al., 2008), protein binding intensity (ChIP-seq) (Robertson et al., 2007), chromatin accessibility (DHS and ATAC-seq) (Buenrostro et al., 2015), and protein-mediated long-range chromatin interactions (ChIA-PET) (Fullwood et al., 2009). Members of the ENCODE consortium (ENCODE Project Consortium, 2012) have collectively generated these datasets in diverse organisms, tissues, and cell types. The 1-dimensional (1-D) signal tracks of the datasets are generally stored in a bigWig compressed binary format or in a bedGraph text format. Although bigWig is a space-efficient standard format for visualizing data on genome browsers, the bedGraph format is often used for text processing and downstream analyses.

A common task in analyzing 1-D signals is extracting summary statistics of a given genomic region. For instance, it is useful to compare an average binding intensity in a peak region of the ChIP-seq signal track to that in a non-peak region. When analyzing new assays with unknown background null distributions, one may need to randomly sample as many as 10 billion regions to obtain sufficient statistical power to assess the significance of observed data for de-noising (Zheng et al., 2019). Thus, a fast algorithm is highly desirable. To accommodate this feature in the widely used Python language, we developed a package pyBedGraph and demonstrate its ability to quickly obtain summary statistics directly from a bedGraph file without the need to convert it to bigWig. The features of pyBedGraph include finding: 1) exact mean, minimum, maximum, coverage, and standard deviations; 2) approximate solutions to the mean.

## 2 Methods

Searching for a given interval in a large bedGraph file is a computationally expensive job. To overcome this problem, pyBedGraph creates an array that contains an index to an entry of data corresponding to a bedGraph line for every base pair in a chromosome. Therefore, when searching for a statistic, pyBedGraph can then simply use the array indices to rapidly access the bedGraph values, thereby avoiding the need to search.

In addition to finding the exact mean, pyBedGraph offers the option to approximate it with a reduced calculation time. The program can pre-calculate and store bins containing values over non-overlapping windows to substantially decrease the number of values indexed and hence the runtime. In this method, pyBedGraph looks up the two bins containing the start and end of the interval and inclusively extracts all bins between the two. When the first and last bin do not exactly match the start and end of the interval, respectively, an estimate is made for each bin by taking the (value of the bin) × (proportion of the bin overlapping the interval). This method trades off the speed with accuracy.

pyBedGraph is implemented in Python3 using Cython to further optimize speed. Detailed methods are provided in **Supplementary data**.

## 3 Results

We benchmarked the performance of pyBedGraph and its bigWig counterpart pyBigWig (Ramírez *et al*., 2016) on 6 ChIP-seq and 2 ATAC-seq mammalian datasets (**Supplementary data**) downloaded from the ENCODE portal (Sloan *et al*., 2016) (https://www.encodeproject.org). All runs were on a Intel(R) Core(TM) i5-7300HQ CPU @ 2.50GHz with 16 GB of RAM using a single thread.

### 3.1 Speed

Using an interval size of 500 bp and bin sizes of 100, 50, or 25 bp, we measured the runtime of looking up 0.1 to 1 million intervals from chr1. The results are illustrated for POLR2A ChIP-seq data (‘ENCFF376VCU’), where pyBedGraph takes 0.26 second (cf. 56 seconds for pyBigWig) to obtain an exact mean in 1 million intervals (**Figure 1a**). Our approximate computation takes 0.09, 0.11, and 0.14 seconds for bin sizes 100, 500, and 25 bp, respectively, while pyBigWig takes 56 seconds. As the size of the query intervals get larger, the run time gradually decreases for pyBedGraph’s approximate mean while it increases for the calculation of the exact mean (**Supplementary data**).

**Fig. 1.**
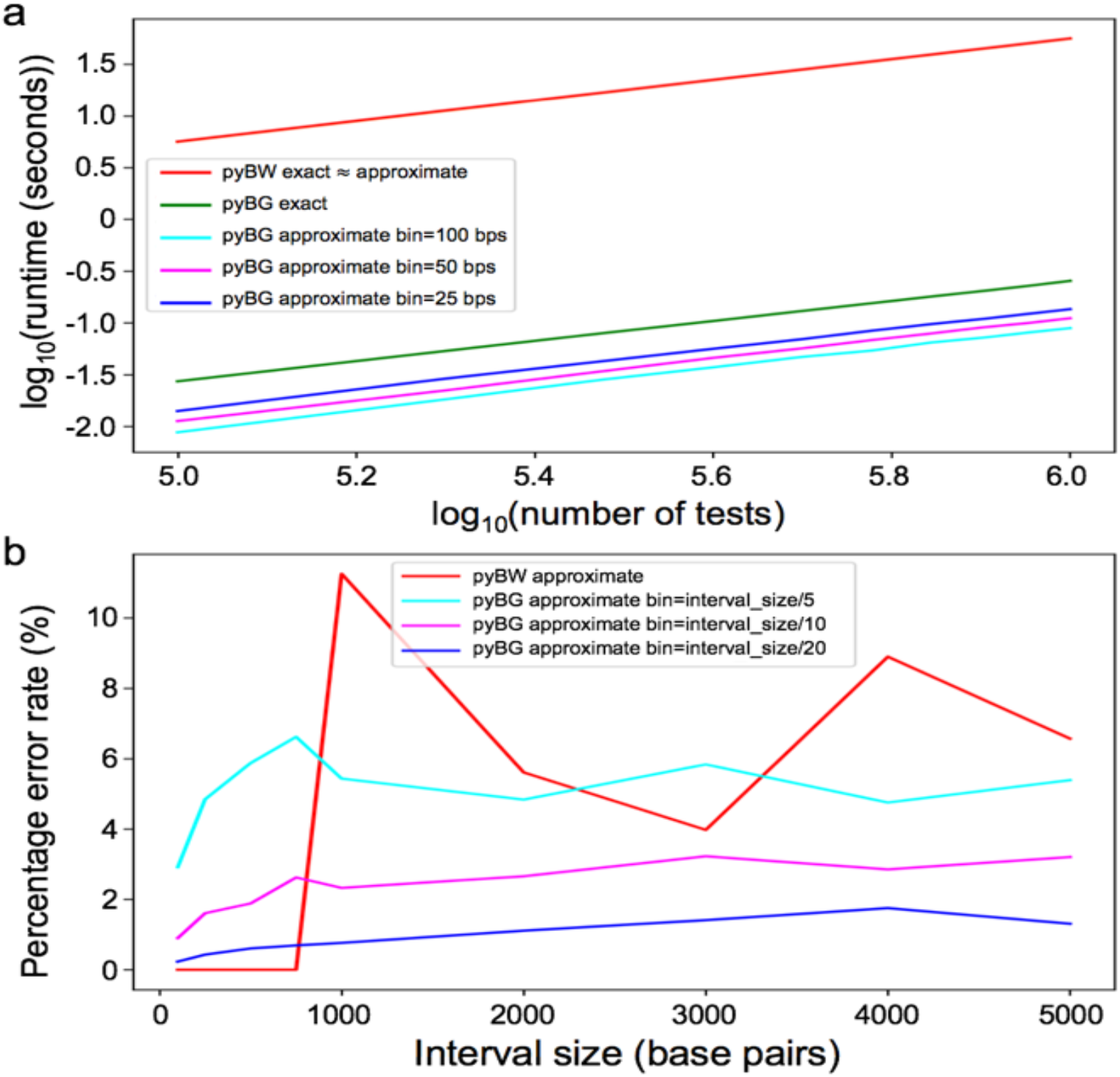
Speed and accuracy benchmark on ENCFF376VCU dataset. a) Runtimes of `pyBigWig` (pyBW) and `pyBedGraph` (pyBG) are recorded for 0.1 to 1 million intervals of size 500 bps. The approximate algorithm for pyBG uses bin sizes of 100, 50, 25 bps. b) The percentage error rate is calculated for approximate solutions as a function of interval sizes ranging from 100 bps to 5000 bps, each with 10,000 intervals to test. For pyBG, bin sizes are the interval size divided by 5, 10, and 20.

### 3.2 Accuracy

We next measured the amount of error resulting from the approximation. For each interval size from 100 bp to 5,000 bp, the percentage error was defined as 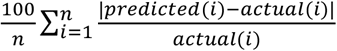, where *n* = 10,000 is the number of regions to look up (test case intervals) in chr1. A test case interval was excluded from the error calculation when its actual value was ‘None’ or 0 while the predicted was not, occurring in less than 2.5 percent of test cases. Mean squared errors and absolute errors were also computed (**Supplementary data**). On the ‘ENCFF376VCU’ dataset, the error was around 6%, 3%, and 1% for pyBedGraph with bin sizes equal to the interval size divided by 5, 10, and 20, respectively (**Figure 1b**). By contrast, pyBigWig utilizes ‘zoom levels’ in the bigWig file and its approximation error peaked at 11% and 9% for interval sizes of 1,000 bps and 4,000 bps, respectively.

### 3.2 Memory

The memory usage of pyBedGraph depends on the number of lines in the bedGraph file and sizes of each chromosome in the reference genome. Reading in the bedGraph file stores two 32-bit integers and a 64-bit float for each line; “loading” each chromosome creates an array of 32-bit integers to store the location of the corresponding line from the bedGraph file. Furthermore, the pre-calculated bins are stored in an array of 64-bit floats of length equal to the genome size divided by the bin size. For example, a bedGraph file for POLR2A ChIP-seq data in mouse spleen (‘ENCFF376VCU’) uses ~1.6GB (=100,620,515 lines × (4 + 4 + 8) bytes) of memory to load the whole genome and an additional 0.8GB (= 2.0 × 10^8^ base pair × 4 bytes) to load chromosome 1 (chr1). Storing bins of size 100 base pairs (bps) requires only 16MB 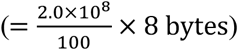. The memory usage of pyBedGraph is nontrivial, yet still reasonable for most laptops. By contrast, pyBigWig does not load the bigwig file and consequently uses no memory.

## 4 Discussion

We developed pyBedGraph and demonstrated its ability to quickly obtain summary statistics from 1-dimensional genomic signals in bedGraph format. Specifically, obtaining the exact mean for 10 billion intervals is estimated to take 43 minutes with pyBedGraph and 7.4 days with pyBigWig. However, one drawback of pyBedGraph is that it can take up to a minute to load files whereas pyBigWig allows instant computation. Therefore, we recommend users to choose pyBedGraph if they prefer working with bedGraph file instead of bigWig, or if they have to search within more than 1 million intervals. For more than 1 billion intervals with limited compute time, our approximate solution with a small bin size may be a viable option. As genomics researchers continue to develop novel technologies ranging from bulk cells to single-cell and single-molecule experiments, it will be imperative to distinguish true signal from technical noise. Particularly, some ChIP-seq, ChIA-PET, and ChIA-Drop experiments yield only 10-20% enrichment rates due to weak antibody, resulting in noisy tracks. We envision pyBedGraph to play a vital role in quickly sampling null distributions to help researchers to de-noise the data.

## Supplementary Method, Tables and Figures

**Supplementary Method S1.** Example mean statistics and error calculation

**Supplementary Method S2.** `pyBedGraph` usage and commands

**Supplementary Method S1**. Example mean statistics and error calculation

An example bedGraph file is given by the following:

|  |  |  |  |
| --- | --- | --- | --- |
| chr1 | 0 | 2 | 0.7 |
| chr1 | 2 | 5 | 0.9 |
| chr1 | 8 | 9 | 0.1 |
| chr1 | 13 | 14 | 0.9 |
| chr1 | 19 | 20 | 0.4 |

After reading the file, the bedGraph object stores the following table as three NumPy arrays for ‘Start’, ‘End’, and ‘Value’.

|  |  |  |  |  |  |
| --- | --- | --- | --- | --- | --- |
| Index | 0 | 1 | 2 | 3 | 4 |
| Start | 0 | 2 | 8 | 13 | 19 |
| End | 2 | 5 | 9 | 14 | 20 |
| Value | 0.7 | 0.9 | 0.1 | 0.9 | 0.4 |

Loading “chr1” creates the following array used to skip the searching process.

|  |  |  |  |  |  |  |  |  |  |  |  |  |  |  |  |  |  |  |  |  |
| --- | --- | --- | --- | --- | --- | --- | --- | --- | --- | --- | --- | --- | --- | --- | --- | --- | --- | --- | --- | --- |
| Index | 0 | 1 | 2 | 3 | 4 | 5 | 6 | 7 | 8 | 9 | 10 | 11 | 12 | 13 | 14 | 15 | 16 | 17 | 18 | 19 |
| Value | 0 | 0 | 1 | 1 | 1 | -1 | -1 | -1 | 2 | -1 | -1 | -1 | -1 | 3 | -1 | -1 | -1 | -1 | -1 | 4 |

**Exact statistics:**

If the user searches the interval [‘chr1’, 8, 12], `pyBedGraph` begins by looking up the 8^th^ index and obtains the value ‘2’. It then goes to the 2^nd^ index in the arrays stored in the bedGraph object. The value ‘0.1’ and length of the interval (9 – 8 = 1) at the 2^nd^ index is used to calculate the relevant statistic. Moving on to the next interval at the 3^rd^ index, `pyBedGraph` sees that the “Start” (‘13’) is outside our query interval and stops looking further.

**Approximate mean:**

If bins of size 5 are used for storing the above values, the following bin array is created:

|  |  |  |  |  |
| --- | --- | --- | --- | --- |
| Index | 0 | 1 | 2 | 3 |
| Value | $2 * 0.7 + 3 * 0.9 = 4.1$ | $1 * 0.1 = 0.1$ | $1 * 0.9 = 0.9$ | $1 * 0.4 = 0.4$ |

If the user searches the interval [‘chr1’, 0, 6], `pyBedGraph` notes that the starting index in the bin array is 0 / 5 = 0 and the end index is floor(6/5) = floor(1.2) = 1. Since the 0^th^ bin is completely inside the interval, the value is stored with the weight of 5 (=bin size). The next bin with index 1 is only partly in the interval so the value is stored with the weight of 1 (=6 mod 5). The calculation found is then (4.1 + 0.1 / 5) / (5 + 5 * 1/5) = 0.69 The correct exact mean is (0.7×2 + 0.9×3) / (2 + 3) = 0.82. As a result, the percentage error in this example is 100*|0.69 – 0.82| / 0.82 = 15.9%.

**Supplementary Method S2.** pyBedGraph usage and commands

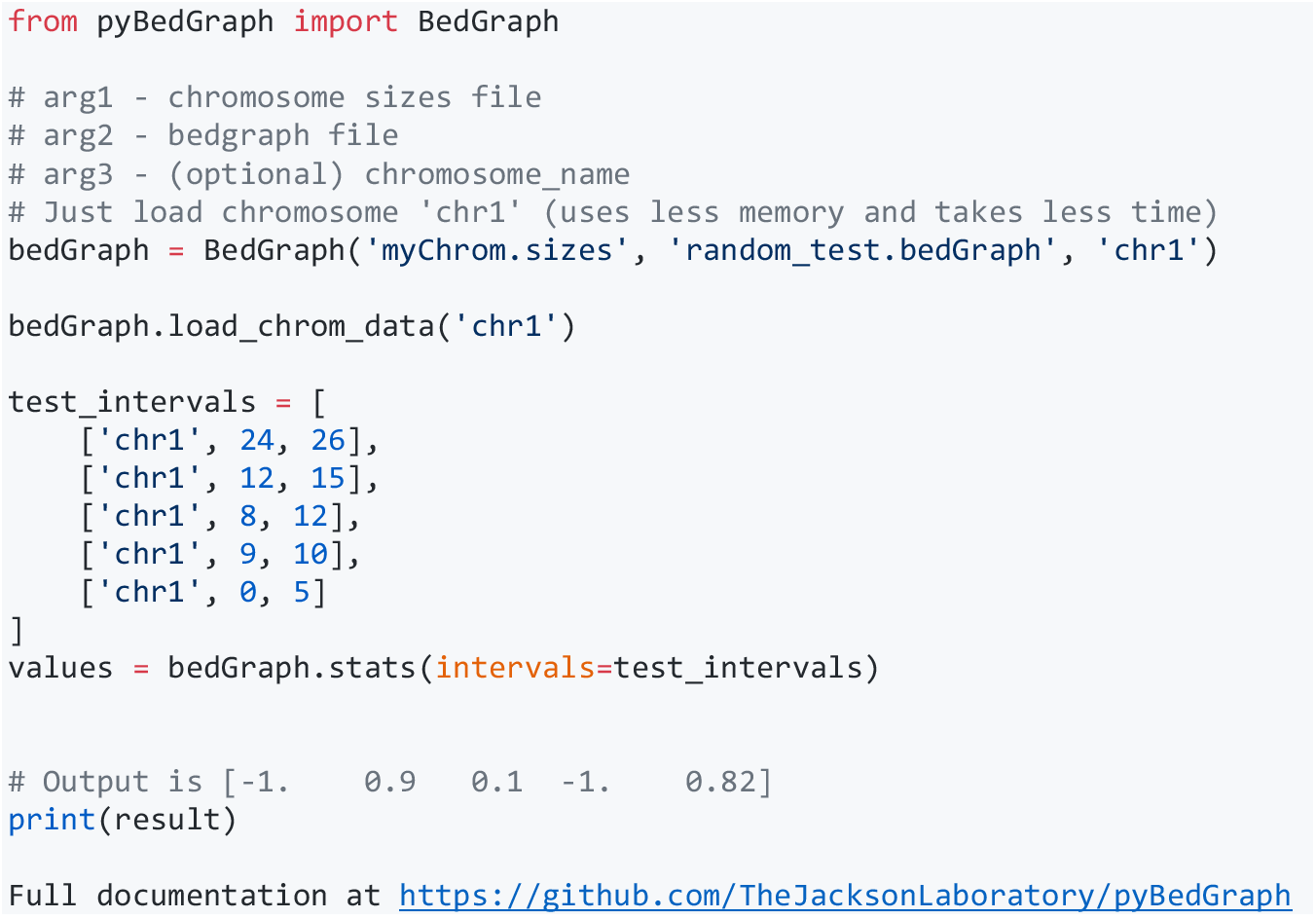

**Supplementary Table S1.** Details of 8 datasets used in the benchmark

| ID | Assay type | Factor | Organism | Genome assembly | Cell/tissue type |
| --- | --- | --- | --- | --- | --- |
| ENCFF376VCU | ChIP-seq | POLR2A | *Mus musculus* C57BL/6 | mm10 | Adult spleen |
| ENCFF643WMY | ChIP-seq | ZNF384 | *Mus musculus* 129S | mm10 | ES-E14 |
| ENCFF384CMP | ChIP-seq | H3K9me3 | *Mus musculus* C57BL/6 | mm10 | Forebrain embryo |
| ENCFF321FZQ | ChIP-seq | POLR2A | *Homo sapiens* | GRCh38 | K562 |
| ENCFF050CCI | ChIP-seq | CTCF | *Homo sapiens* | GRCh38 | K562 |
| ENCFF847JMY | ChIP-seq | H3K4me3 | *Homo sapiens* | GRCh38 | K562 |
| ENCFF631HEX | ATAC-seq | N/A | *Homo sapiens* | GRCh38 | A549 |
| ENCFF770CQD | ATAC-seq | N/A | *Mus musculus* C57BL/6 | mm10 | p0 liver |

The 8 ENCODE datasets used in the benchmark. ID: unique identification assigned by ENCODE; Assay type: experimental assay; Factor: protein immunoprecipitation factor if ChIP-seq; Organism: name of the species and strain (if *Mus musculus*); Genome assembly: reference genome used for mapping reads; Cell/tissue type: name of the cell line or tissue as listed by ENCODE.

**Supplementary Table S2.** Runtime for 1 million test intervals

| Dataset | pyBW exact | pyBW app. | pyBG exact | pyBG app.<br>Bin=100 bps | pyBG app.<br>Bin=50 bps | pyBG app.<br>Bin=25 bps |
| --- | --- | --- | --- | --- | --- | --- |
| ENCFF050CCI | 59.265 | 59.649 | 0.266 | 0.093 | 0.113 | 0.137 |
| ENCFF321FZQ | 62.871 | 126.587 | 0.373 | 0.092 | 0.115 | 0.139 |
| ENCFF376VCU | 55.952 | 56.134 | 0.254 | 0.089 | 0.111 | 0.135 |
| ENCFF384CMP | 54.093 | 54.245 | 0.252 | 0.089 | 0.111 | 0.136 |
| ENCFF631HEX | 117.052 | 117.129 | 0.216 | 0.093 | 0.115 | 0.139 |
| ENCFF643WMY | 55.591 | 55.774 | 0.257 | 0.090 | 0.112 | 0.137 |
| ENCFF770CQD | 52.306 | 52.282 | 0.255 | 0.089 | 0.111 | 0.136 |
| ENCFF847JMY | 56.080 | 56.408 | 0.215 | 0.092 | 0.115 | 0.137 |
| AVERAGE | 64.151 | 72.276 | 0.261 | 0.091 | 0.113 | 0.137 |

Runtime recorded in seconds, for all 8 datasets. pyBW: pyBigWig; pyBG: pyBedGraph; app.: approximation.

**Supplementary Table S3.** Error statistics for ENCFF050CCI and ENCFF321FZQ

| **a** ENCFF050CCI --- pyBW app. |  |  |  |  | **b** ENCFF050CCI --- pyBG app. bin=int_size/20 |  |  |  |  |
| --- | --- | --- | --- | --- | --- | --- | --- | --- | --- |
| Interval Size (bPS) | Error Rate (%) | Mean Squared Error | Absolute Error | # Actual is 0 | Interval Size (bPS) | Error Rate (%) | Mean Squared Error | Absolute Error | # Actual is 0 |
| 100 | 0.0 | 0.0 | 0.0 | 0 | 100 | 0.19777 | 0.0 | 0.00051 | 25 |
| 250 | 0.0 | 0.0 | 0.0 | 0 | 250 | 0.44277 | 1e-05 | 0.001 | 18 |
| 500 | 0.0 | 0.0 | 0.0 | 0 | 500 | 0.5064 | 2e-05 | 0.00172 | 14 |
| 750 | 0.0 | 0.0 | 0.0 | 0 | 750 | 0.43736 | 4e-05 | 0.0024 | 7 |
| 1000 | 8.84954 | 0.04867 | 0.05697 | 59 | 1000 | 0.56813 | 6e-05 | 0.0027 | 6 |
| 2000 | 4.86489 | 0.01083 | 0.0265 | 29 | 2000 | 0.8495 | 0.00023 | 0.00427 | 8 |
| 3000 | 2.73314 | 0.00566 | 0.01925 | 16 | 3000 | 0.83133 | 0.00043 | 0.0056 | 3 |
| 4000 | 7.37763 | 0.02792 | 0.05276 | 51 | 4000 | 0.87903 | 0.00051 | 0.00599 | 10 |
| 5000 | 6.096 | 0.0217 | 0.04282 | 39 | 5000 | 1.0037 | 0.00091 | 0.00701 | 9 |

| **c** ENCFF050CCI --- pyBG app. bin=int_size/5 |  |  |  |  | **d** ENCFF050CCI --- pyBG app. bin=int_size/10 |  |  |  |  |
| --- | --- | --- | --- | --- | --- | --- | --- | --- | --- |
| Interval Size (bPS) | Error Rate (%) | Mean Squared Error | Absolute Error | # Actual is 0 | Interval Size (bPS) | Error Rate (%) | Mean Squared Error | Absolute Error | # Actual is 0 |
| 100 | 2.22438 | 0.00055 | 0.00604 | 112 | 100 | 0.69011 | 4e-05 | 0.00182 | 62 |
| 250 | 4.20873 | 0.00139 | 0.01108 | 79 | 250 | 1.35438 | 0.00016 | 0.00352 | 42 |
| 500 | 3.76417 | 0.00325 | 0.01698 | 48 | 500 | 1.42765 | 0.00027 | 0.00552 | 23 |
| 750 | 4.33152 | 0.00671 | 0.02188 | 27 | 750 | 1.41773 | 0.00066 | 0.00729 | 10 |
| 1000 | 3.94969 | 0.01006 | 0.02452 | 33 | 1000 | 1.51551 | 0.00116 | 0.00852 | 19 |
| 2000 | 4.86489 | 0.01083 | 0.0265 | 29 | 2000 | 2.45116 | 0.00216 | 0.01197 | 16 |
| 3000 | 3.84557 | 0.0131 | 0.02781 | 22 | 3000 | 1.91895 | 0.00311 | 0.01359 | 12 |
| 4000 | 3.77184 | 0.0088 | 0.02642 | 29 | 4000 | 1.9365 | 0.00306 | 0.01362 | 17 |
| 5000 | 4.00455 | 0.0101 | 0.02794 | 28 | 5000 | 2.17891 | 0.00366 | 0.01504 | 14 |

**ENCFF321FZQ**
| **e** ENCFF321FZQ --- pyBW app. |  |  |  |  | **f** ENCFF321FZQ --- pyBG app. bin=int_size/20 |  |  |  |  |
| --- | --- | --- | --- | --- | --- | --- | --- | --- | --- |
| Interval Size (bPS) | Error Rate (%) | Mean Squared Error | Absolute Error | # Actual is 0 | Interval Size (bPS) | Error Rate (%) | Mean Squared Error | Absolute Error | # Actual is 0 |
| 100 | 0.0 | 0.0 | 0.0 | 0 | 100 | 0.05095 | 0.0 | 0.00018 | 14 |
| 250 | 0.0 | 0.0 | 0.0 | 0 | 250 | 0.18585 | 0.0 | 0.00035 | 18 |
| 500 | 5.80243 | 0.00441 | 0.01406 | 59 | 500 | 0.24962 | 1e-05 | 0.00062 | 14 |
| 750 | 2.79372 | 0.00942 | 0.01072 | 20 | 750 | 0.22616 | 9e-05 | 0.00089 | 3 |
| 1000 | 1.68375 | 0.00096 | 0.00746 | 13 | 1000 | 0.23464 | 1e-05 | 0.001 | 2 |
| 2000 | 4.93825 | 0.00941 | 0.02458 | 13 | 2000 | 0.31463 | 2e-05 | 0.00152 | 2 |
| 3000 | 3.32866 | 0.00246 | 0.01662 | 9 | 3000 | 0.40396 | 7e-05 | 0.00206 | 3 |
| 4000 | 2.75904 | 0.0013 | 0.01222 | 8 | 4000 | 0.51945 | 0.0001 | 0.00241 | 3 |
| 5000 | 2.14842 | 0.00131 | 0.00997 | 6 | 5000 | 0.58974 | 0.00018 | 0.00293 | 0 |

| **g** ENCFF321FZQ --- pyBG app. bin=int_size/5 |  |  |  |  | **h** ENCFF321FZQ --- pyBG app. bin=int_size/10 |  |  |  |  |
| --- | --- | --- | --- | --- | --- | --- | --- | --- | --- |
| Interval Size (bPS) | Error Rate (%) | Mean Squared Error | Absolute Error | # Actual is 0 | Interval Size (bPS) | Error Rate (%) | Mean Squared Error | Absolute Error | # Actual is 0 |
| 100 | 0.78766 | 2e-05 | 0.00198 | 72 | 100 | 0.25667 | 0.0 | 0.00063 | 34 |
| 250 | 2.1133 | 0.0001 | 0.00402 | 75 | 250 | 0.70887 | 1e-05 | 0.00119 | 42 |
| 500 | 2.28113 | 0.00048 | 0.00609 | 46 | 500 | 0.79472 | 0.00011 | 0.00198 | 26 |
| 750 | 2.52241 | 0.00187 | 0.00866 | 16 | 750 | 0.69359 | 0.00018 | 0.00262 | 10 |
| 1000 | 2.29411 | 0.00313 | 0.01034 | 22 | 1000 | 0.75259 | 9e-05 | 0.00313 | 9 |
| 2000 | 2.75565 | 0.00294 | 0.01398 | 13 | 2000 | 0.99123 | 0.00105 | 0.00512 | 8 |
| 3000 | 2.90612 | 0.00172 | 0.01399 | 13 | 3000 | 1.21472 | 0.00064 | 0.00629 | 7 |
| 4000 | 3.36092 | 0.00319 | 0.01499 | 8 | 4000 | 1.53006 | 0.0007 | 0.00698 | 6 |
| 5000 | 2.70569 | 0.00255 | 0.01477 | 11 | 5000 | 1.4146 | 0.00081 | 0.00737 | 3 |

Additional error statistics for approximate results from pyBigWig and pyBedGraph. Mean squared error is calculated as $(actual - predicted)^2$ , and the absolute error is calculated as $|actual - predicted|$ . “# actual is 0” is the number of test cases not included in the percentage error rate due to the actual result being 0 while predicted was not. ENCFF050CCI: CTCF ChIP-seq in human K562 cell line; ENCFF321FZQ: POLR2A ChIP-seq in human K562 cell line. Tables S4, S5, S6 follow the same notation as Table S3.

**Supplementary Table S4.** Error statistics for ENCFF631HEX and ENCFF847JMY

ENCFF631HEX
| a ENCFF631HEX --- pyBW app. |  |  |  |  | b ENCFF631HEX --- pyBG app. bin=int_size/20 |  |  |  |  |
| --- | --- | --- | --- | --- | --- | --- | --- | --- | --- |
| Interval Size (bPS) | Error Rate (%) | Mean Squared Error | Absolute Error | # Actual is 0 | Interval Size (bPS) | Error Rate (%) | Mean Squared Error | Absolute Error | # Actual is 0 |
| 100 | 0.0 | 0.0 | 0.0 | 0 | 100 | 0.15783 | 0.00741 | 0.02161 | 25 |
| 250 | 0.0 | 0.0 | 0.0 | 0 | 250 | 0.32185 | 0.01453 | 0.04444 | 46 |
| 500 | 0.0 | 0.0 | 0.0 | 0 | 500 | 0.38188 | 0.03805 | 0.08614 | 37 |
| 750 | 0.0 | 0.0 | 0.0 | 0 | 750 | 0.46332 | 0.0574 | 0.1167 | 26 |
| 1000 | 0.0 | 0.0 | 0.0 | 0 | 1000 | 0.52255 | 0.08775 | 0.14388 | 17 |
| 2000 | 12.87664 | 231.13895 | 5.64724 | 47 | 2000 | 0.60204 | 0.22035 | 0.23684 | 5 |
| 3000 | 7.7858 | 97.29978 | 3.81412 | 35 | 3000 | 0.64027 | 0.37413 | 0.30447 | 11 |
| 4000 | 5.23807 | 54.82365 | 2.81183 | 20 | 4000 | 0.77547 | 0.63408 | 0.3697 | 4 |
| 5000 | 4.85157 | 32.97427 | 2.26198 | 11 | 5000 | 0.99399 | 0.73966 | 0.41869 | 3 |

| c ENCFF631HEX --- pyBG app. bin=int_size/5 |  |  |  |  | d ENCFF631HEX --- pyBG app. bin=int_size/10 |  |  |  |  |
| --- | --- | --- | --- | --- | --- | --- | --- | --- | --- |
| Interval Size (bPS) | Error Rate (%) | Mean Squared Error | Absolute Error | # Actual is 0 | Interval Size (bPS) | Error Rate (%) | Mean Squared Error | Absolute Error | # Actual is 0 |
| 100 | 1.60011 | 0.47989 | 0.28938 | 125 | 100 | 0.52078 | 0.06076 | 0.08329 | 59 |
| 250 | 3.20007 | 1.46303 | 0.5661 | 175 | 250 | 1.10833 | 0.1454 | 0.1676 | 89 |
| 500 | 3.93499 | 3.456 | 0.91131 | 140 | 500 | 1.29715 | 0.35248 | 0.28417 | 76 |
| 750 | 4.14189 | 6.84428 | 1.20181 | 108 | 750 | 1.60001 | 0.58176 | 0.38092 | 57 |
| 1000 | 4.41435 | 9.50925 | 1.43425 | 63 | 1000 | 1.45005 | 0.89312 | 0.46396 | 37 |
| 2000 | 6.01991 | 18.63036 | 1.96445 | 25 | 2000 | 1.74513 | 2.36786 | 0.73055 | 13 |
| 3000 | 4.2207 | 26.81737 | 2.16785 | 29 | 3000 | 1.93702 | 4.20412 | 0.91446 | 17 |
| 4000 | 4.8659 | 33.5904 | 2.23606 | 18 | 4000 | 1.96842 | 4.91793 | 0.99207 | 9 |
| 5000 | 4.4646 | 35.01066 | 2.26956 | 11 | 5000 | 2.30428 | 5.66221 | 1.0355 | 9 |

ENCFF847JMY
| e ENCFF847JMY --- pyBW app. |  |  |  |  | f ENCFF847JMY --- pyBG app. bin=int_size/20 |  |  |  |  |
| --- | --- | --- | --- | --- | --- | --- | --- | --- | --- |
| Interval Size (bPS) | Error Rate (%) | Mean Squared Error | Absolute Error | # Actual is 0 | Interval Size (bPS) | Error Rate (%) | Mean Squared Error | Absolute Error | # Actual is 0 |
| 100 | 0.0 | 0.0 | 0.0 | 0 | 100 | 0.16888 | 0.0 | 0.00025 | 38 |
| 250 | 0.0 | 0.0 | 0.0 | 0 | 250 | 0.61892 | 1e-05 | 0.00056 | 60 |
| 500 | 0.0 | 0.0 | 0.0 | 0 | 500 | 1.12473 | 4e-05 | 0.00113 | 75 |
| 750 | 0.0 | 0.0 | 0.0 | 0 | 750 | 1.06239 | 7e-05 | 0.00155 | 44 |
| 1000 | 0.0 | 0.0 | 0.0 | 0 | 1000 | 0.91102 | 0.00014 | 0.0021 | 28 |
| 2000 | 13.91072 | 0.18049 | 0.07293 | 73 | 2000 | 0.8792 | 0.00028 | 0.00343 | 14 |
| 3000 | 7.9812 | 0.06931 | 0.04792 | 42 | 3000 | 1.02717 | 0.00039 | 0.00437 | 9 |
| 4000 | 6.79703 | 0.0409 | 0.03638 | 35 | 4000 | 1.63952 | 0.00063 | 0.00514 | 10 |
| 5000 | 4.96046 | 0.01967 | 0.02691 | 27 | 5000 | 1.2845 | 0.00088 | 0.00576 | 4 |

| g ENCFF847JMY --- pyBG app. bin=int_size/5 |  |  |  |  | h ENCFF847JMY --- pyBG app. bin=int_size/10 |  |  |  |  |
| --- | --- | --- | --- | --- | --- | --- | --- | --- | --- |
| Interval Size (bPS) | Error Rate (%) | Mean Squared Error | Absolute Error | # Actual is 0 | Interval Size (bPS) | Error Rate (%) | Mean Squared Error | Absolute Error | # Actual is 0 |
| 100 | 2.24585 | 0.00035 | 0.00352 | 220 | 100 | 0.7316 | 3e-05 | 0.00096 | 99 |
| 250 | 5.77551 | 0.00182 | 0.00751 | 307 | 250 | 1.98588 | 0.00015 | 0.00215 | 152 |
| 500 | 9.68363 | 0.00362 | 0.01231 | 242 | 500 | 3.53039 | 0.00045 | 0.00392 | 138 |
| 750 | 9.34833 | 0.00889 | 0.01684 | 132 | 750 | 3.63639 | 0.00091 | 0.00536 | 73 |
| 1000 | 8.41881 | 0.01237 | 0.02097 | 96 | 1000 | 2.99831 | 0.00134 | 0.00681 | 51 |
| 2000 | 6.03585 | 0.02269 | 0.0256 | 34 | 2000 | 2.75019 | 0.0028 | 0.0105 | 21 |
| 3000 | 5.15734 | 0.02357 | 0.02715 | 33 | 3000 | 2.58095 | 0.00384 | 0.01199 | 17 |
| 4000 | 6.44138 | 0.02428 | 0.02795 | 35 | 4000 | 4.44111 | 0.00468 | 0.01249 | 20 |
| 5000 | 4.94017 | 0.02465 | 0.02893 | 26 | 5000 | 2.53949 | 0.00415 | 0.01258 | 11 |

Refer to Table S3 legend for labels. ENCFF631HEX: ATAC-seq in human A549; ENCFF847JMY: H3K4me3 ChIP-seq in human K562 cell line.

**Supplementary Table S5.**
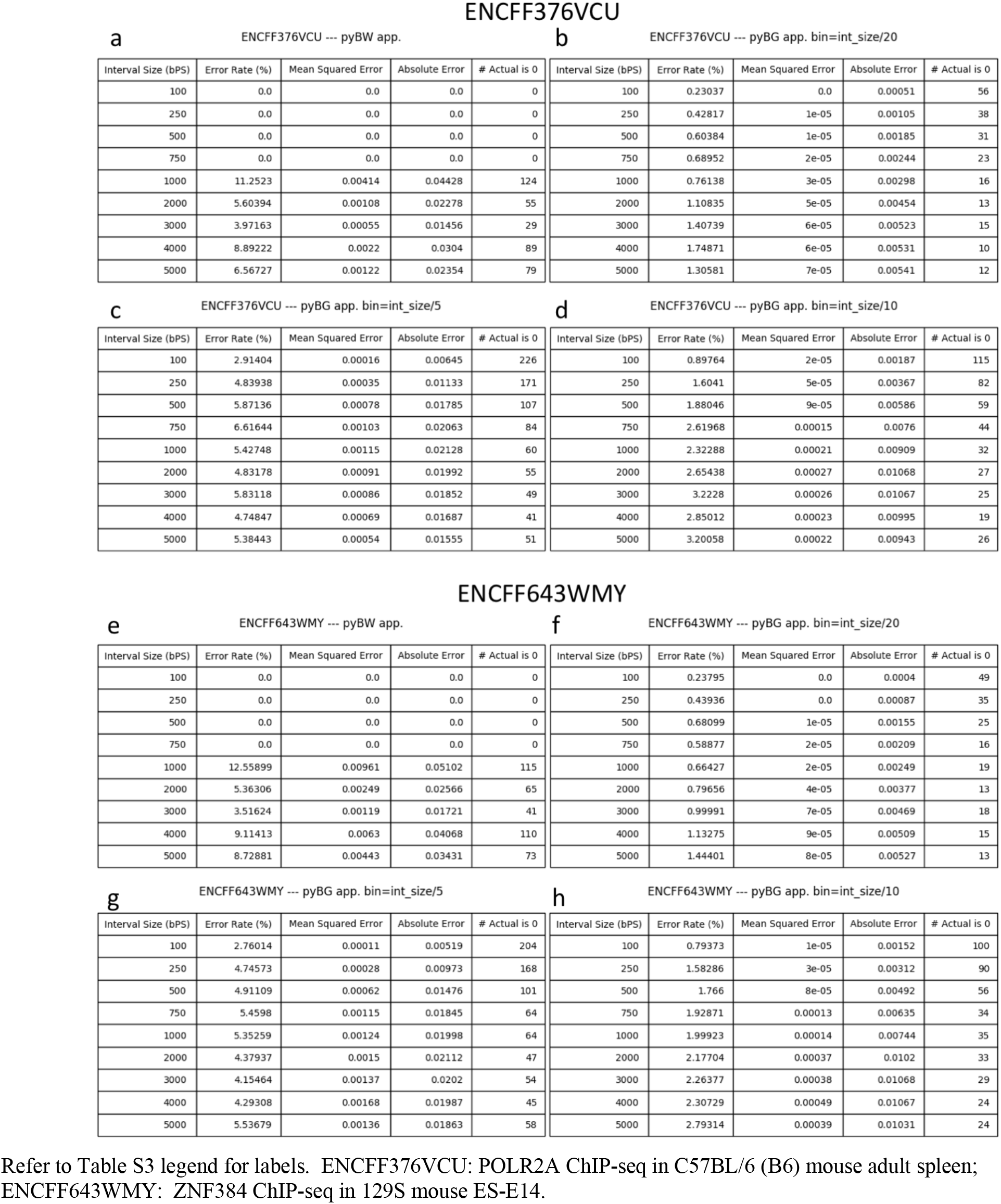
Error statistics for ENCFF376VCU and ENCFF643WMY

**Supplementary Table S6.** Error statistics for ENCFF384CMP and ENCFF770CQD

| **a** ENCFF384CMP --- pyBW app. |  |  |  |  | **b** ENCFF384CMP --- pyBG app. bin=int_size/20 |  |  |  |  |
| --- | --- | --- | --- | --- | --- | --- | --- | --- | --- |
| Interval Size (bPS) | Error Rate (%) | Mean Squared Error | Absolute Error | # Actual is 0 | Interval Size (bPS) | Error Rate (%) | Mean Squared Error | Absolute Error | # Actual is 0 |
| 100 | 0.0 | 0.0 | 0.0 | 0 | 100 | 0.17917 | 0.0 | 0.00038 | 57 |
| 250 | 0.0 | 0.0 | 0.0 | 0 | 250 | 0.48411 | 0.0 | 0.00081 | 63 |
| 500 | 0.0 | 0.0 | 0.0 | 0 | 500 | 0.65117 | 1e-05 | 0.00147 | 39 |
| 750 | 0.0 | 0.0 | 0.0 | 0 | 750 | 0.53995 | 1e-05 | 0.00195 | 23 |
| 1000 | 15.23345 | 0.0061 | 0.05198 | 105 | 1000 | 0.7096 | 2e-05 | 0.00241 | 21 |
| 2000 | 5.88282 | 0.00148 | 0.026 | 47 | 2000 | 0.82115 | 3e-05 | 0.00368 | 9 |
| 3000 | 4.02822 | 0.00073 | 0.01812 | 27 | 3000 | 1.08191 | 5e-05 | 0.00473 | 9 |
| 4000 | 9.92053 | 0.00431 | 0.04258 | 56 | 4000 | 1.25278 | 6e-05 | 0.0053 | 6 |
| 5000 | 8.81868 | 0.00272 | 0.03614 | 45 | 5000 | 1.80034 | 7e-05 | 0.00555 | 8 |

| **c** ENCFF384CMP --- pyBG app. bin=int_size/5 |  |  |  |  | **d** ENCFF384CMP --- pyBG app. bin=int_size/10 |  |  |  |  |
| --- | --- | --- | --- | --- | --- | --- | --- | --- | --- |
| Interval Size (bPS) | Error Rate (%) | Mean Squared Error | Absolute Error | # Actual is 0 | Interval Size (bPS) | Error Rate (%) | Mean Squared Error | Absolute Error | # Actual is 0 |
| 100 | 2.31138 | 0.0001 | 0.00488 | 233 | 100 | 0.70936 | 1e-05 | 0.0014 | 122 |
| 250 | 4.44706 | 0.00026 | 0.00944 | 255 | 250 | 1.47054 | 3e-05 | 0.00291 | 127 |
| 500 | 5.74615 | 0.00055 | 0.01477 | 144 | 500 | 1.8858 | 7e-05 | 0.0048 | 69 |
| 750 | 5.48654 | 0.00079 | 0.01846 | 79 | 750 | 1.88775 | 0.0001 | 0.00626 | 47 |
| 1000 | 5.77311 | 0.001 | 0.02065 | 53 | 1000 | 2.10746 | 0.00013 | 0.00734 | 30 |
| 2000 | 4.98382 | 0.00103 | 0.02142 | 33 | 2000 | 2.42531 | 0.00024 | 0.01032 | 19 |
| 3000 | 4.92644 | 0.00103 | 0.02137 | 32 | 3000 | 2.65572 | 0.00029 | 0.01123 | 20 |
| 4000 | 4.56237 | 0.00096 | 0.02035 | 22 | 4000 | 2.49997 | 0.00027 | 0.01092 | 10 |
| 5000 | 5.98328 | 0.00086 | 0.01937 | 37 | 5000 | 3.0924 | 0.00026 | 0.01082 | 14 |

**ENCFF770CQD**
| **e** ENCFF770CQD --- pyBW app. |  |  |  |  | **f** ENCFF770CQD --- pyBG app. bin=int_size/20 |  |  |  |  |
| --- | --- | --- | --- | --- | --- | --- | --- | --- | --- |
| Interval Size (bPS) | Error Rate (%) | Mean Squared Error | Absolute Error | # Actual is 0 | Interval Size (bPS) | Error Rate (%) | Mean Squared Error | Absolute Error | # Actual is 0 |
| 100 | 0.0 | 0.0 | 0.0 | 0 | 100 | 0.22878 | 0.0 | 0.00037 | 59 |
| 250 | 0.0 | 0.0 | 0.0 | 0 | 250 | 0.49444 | 0.0 | 0.00076 | 30 |
| 500 | 0.0 | 0.0 | 0.0 | 0 | 500 | 0.81446 | 1e-05 | 0.00141 | 20 |
| 750 | 0.0 | 0.0 | 0.0 | 0 | 750 | 0.91214 | 2e-05 | 0.0019 | 20 |
| 1000 | 11.72576 | 0.00594 | 0.03034 | 112 | 1000 | 1.02285 | 3e-05 | 0.00228 | 14 |
| 2000 | 5.42291 | 0.00137 | 0.01458 | 68 | 2000 | 1.4343 | 5e-05 | 0.00336 | 17 |
| 3000 | 3.54555 | 0.00068 | 0.01036 | 40 | 3000 | 1.54412 | 8e-05 | 0.00389 | 11 |
| 4000 | 11.00703 | 0.00615 | 0.0327 | 119 | 4000 | 1.31571 | 9e-05 | 0.00374 | 17 |
| 5000 | 11.25475 | 0.00418 | 0.02548 | 77 | 5000 | 1.52607 | 0.0001 | 0.00396 | 16 |

| **g** ENCFF770CQD --- pyBG app. bin=int_size/5 |  |  |  |  | **h** ENCFF770CQD --- pyBG app. bin=int_size/10 |  |  |  |  |
| --- | --- | --- | --- | --- | --- | --- | --- | --- | --- |
| Interval Size (bPS) | Error Rate (%) | Mean Squared Error | Absolute Error | # Actual is 0 | Interval Size (bPS) | Error Rate (%) | Mean Squared Error | Absolute Error | # Actual is 0 |
| 100 | 3.11766 | 0.00014 | 0.00475 | 216 | 100 | 0.90828 | 1e-05 | 0.00137 | 106 |
| 250 | 5.01282 | 0.00046 | 0.00883 | 144 | 250 | 1.75512 | 5e-05 | 0.00277 | 71 |
| 500 | 5.92168 | 0.00081 | 0.01298 | 70 | 500 | 2.23482 | 0.00011 | 0.00452 | 35 |
| 750 | 5.56093 | 0.00124 | 0.01491 | 64 | 750 | 2.46739 | 0.00015 | 0.0058 | 36 |
| 1000 | 5.59304 | 0.0013 | 0.01458 | 61 | 1000 | 2.71618 | 0.00021 | 0.00658 | 29 |
| 2000 | 5.42291 | 0.00137 | 0.01458 | 68 | 2000 | 2.77676 | 0.00033 | 0.00748 | 35 |
| 3000 | 5.21979 | 0.00163 | 0.01525 | 73 | 3000 | 2.81085 | 0.00041 | 0.00775 | 33 |
| 4000 | 5.21869 | 0.00172 | 0.01565 | 71 | 4000 | 2.5137 | 0.00036 | 0.00749 | 37 |
| 5000 | 5.383 | 0.00172 | 0.01557 | 63 | 5000 | 2.73866 | 0.00041 | 0.00754 | 33 |

Refer to Table S3 legend for labels. ENCFF384CMP: H3K9me3 ChIP-seq in C57BL/6 mouse forebrain embryo; ENCFF770CQD: ATAC-seq in C57BL/6 p0 liver.

**Supplementary Figure S1.**
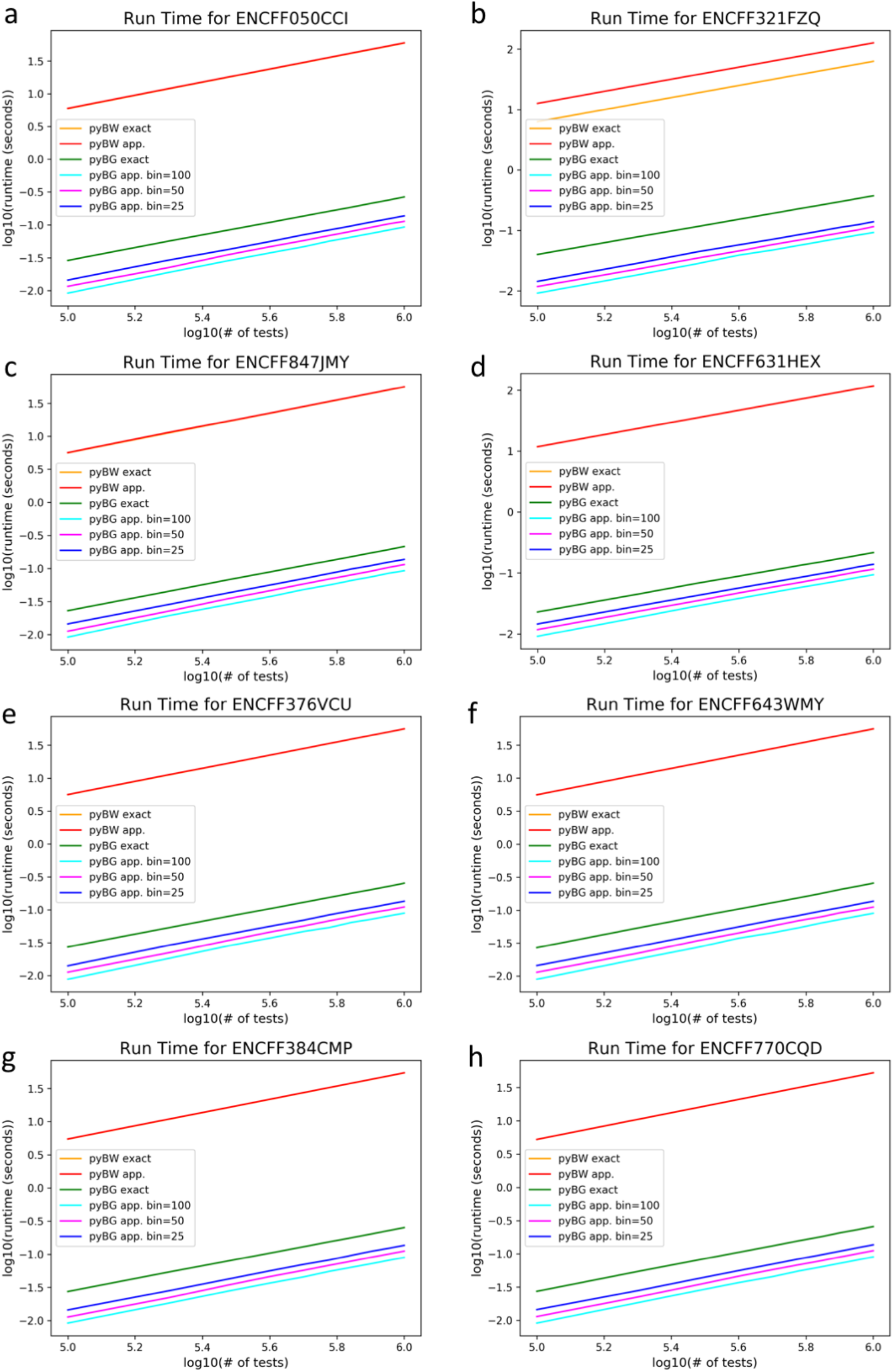
Runtime vs. number of tests for all 8 datasets. Runtime for various numbers of test cases, for all 8 datasets. pyBW: pyBigWig; pyBG: pyBedGraph; app.: approximation. Except for ENCFF321FZQ, pyBW exact has approximately the same runtime as pyBW app., making the red and yellow lines overlap.

**Supplementary Figure S2.**
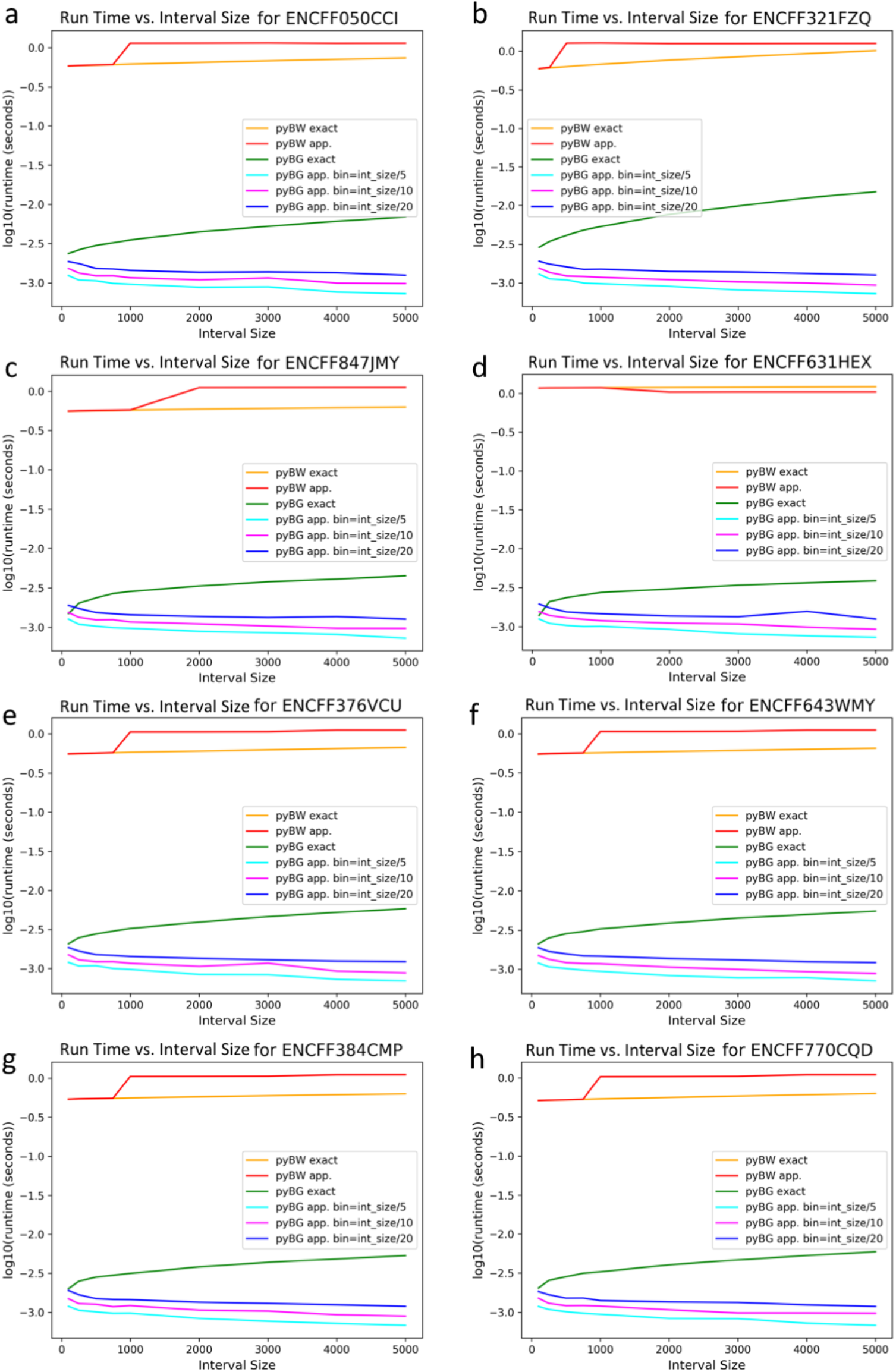
Runtime vs. interval size for all 8 datasets. Runtime for various interval sizes, for all 8 datasets. pyBW: pyBigWig; pyBG: pyBedGraph; app.: approximation.

**Supplementary Figure S3.**
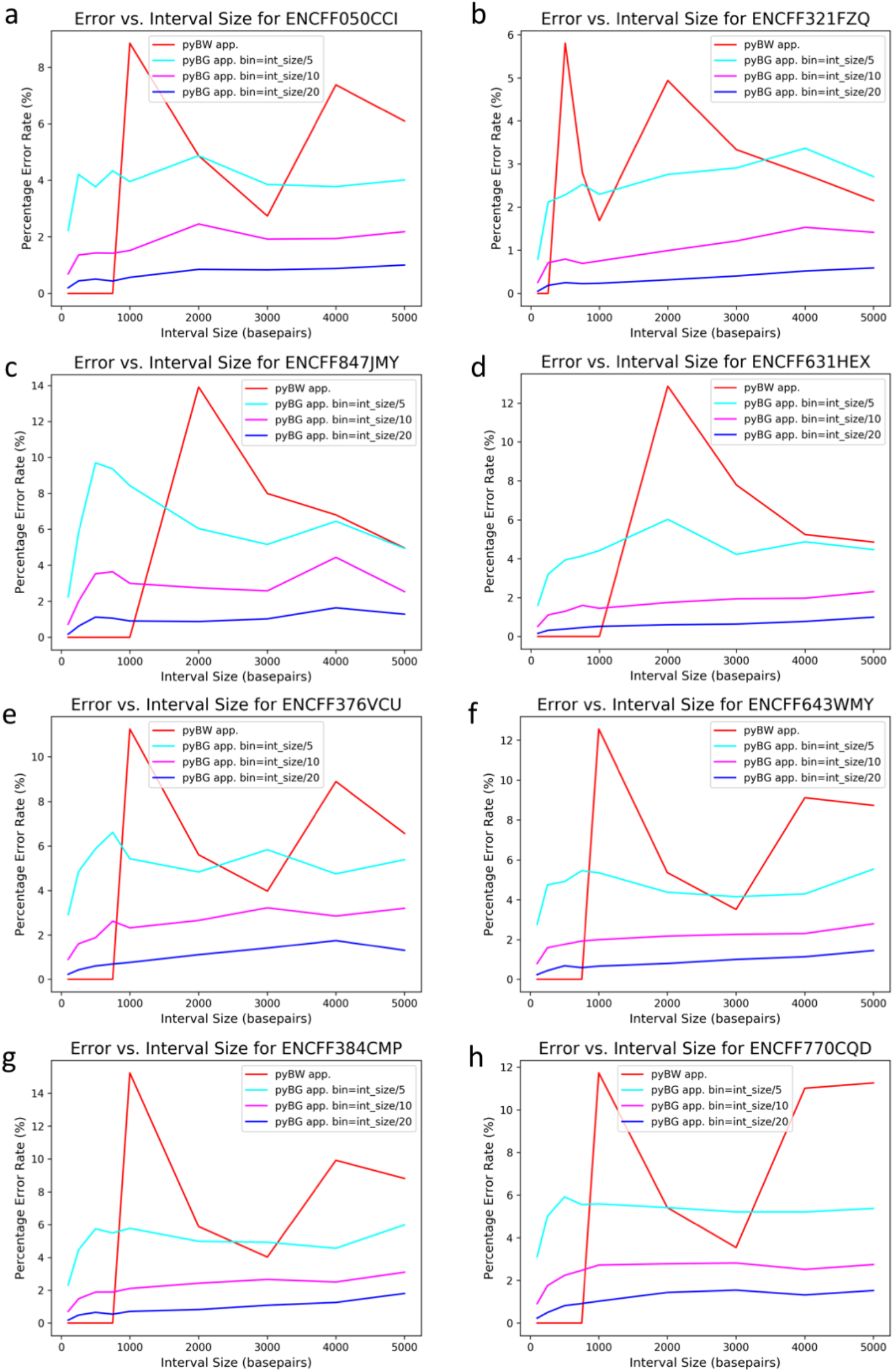
Error rate vs. interval size for all 8 datasets. Percentage error for various interval sizes, for all 8 datasets. pyBW: pyBigWig; pyBG: pyBedGraph; app.: approximation. Errors from exact statistics of both pyBigWig and pyBedGraph are not included since they are zero.

